# Distributed Activity in the Medial Frontal Cortex Predicts Self-Initiated Action

**DOI:** 10.1101/2025.04.01.646546

**Authors:** Julien Claron, Matthieu Blons, Alexandre Dizeux, Thomas Deffieux, Mickael Tanter, Béatrice Berthon, Pierre Pouget

## Abstract

Research indicates that significant damage to primates’ medial frontal cortex (MFC) can impede action initiation when anticipating rewards. However, the specific roles within the MFC related to self-generated action preparation remain debated. Here, while two macaques were engaged in an oculomotor task, they occasionally paused and self-resumed the task after varying amounts of time. We utilized the high signal-to-noise ratio from spatial changes in cerebrovascular blood volume (CBV) recorded with ultrafast ultrasound imaging (fUSi) to determine whether the intervals of breaks or the resumption of task-related CBV signals could predict the execution of an ongoing single action. We show that self-initiated eye movements activate the Supplementary eye field (SEF) and the midcingulate cortex (MCC) up to 5.5 seconds before movement execution. Importantly, our results show that a concurrent activity in the SEF and MCC predicts how these two regions might simultaneously exert opposing influences on the voluntary pause and resume of tasks on a single trial basis. The quantification of this dynamic interaction is a novel step for understanding voluntary action in primates and its ability to initiate one’s movement based on internal motivations rather than driven, instructed behavior.

## INTRODUCTION

Neurophysiological brain responses associated with self-initiated action have been extensively investigated using conventional neurophysiological and imaging techniques^1–3^ (Hallett et al.) In a somewhat controversial experiment, Libet found that subjects reported being aware of their intention to move later than the onset of the electroencephalographic fluctuations (Berenshaft potential) recorded over large medial frontal areas^1^(see also Hallett et al., refs). These results suggest that intentional processes build up during a considerable span of hundreds of milliseconds before the onset of movement. Numerous studies have since explored the subjective feeling of volition, often referred to as “conscious intention” or “motor awareness,” highlighting the crucial roles of the medial frontal cortex (MFC) and pre-supplementary motor area (pre-SMA)^4–10^

Apart from the subjective experiences reported by subjects, various neuroimaging and electrophysiological studies in both humans and non-human primates have investigated the neuronal networks involved in preparing internally generated movements^5,11–16^. Comparisons between free and reflexive movements have similarly identified an extensive fronto-mesial network that includes the pre-SMA and the MCC^12–15^. Paus states that the mesial motor cortex structures and the middle and anterior cingulate cortex are places of convergence for motor control, homeostatic drive, emotion, and cognition^17^.

Several complexities remain in the literature: 1. By definition, self-initiated movements are not experimentally controlled and are consequently infrequent or rare, depending on the subject’s intention. As a result, these isolated events can be complex to analyze due to a low signal-to-noise ratio in methods such as event related potentials (ERPs) and functional magnetic resonance imaging (fMR) (Hallett et al.). 2. The responses may depend on micro-scale and mesoscale neuronal activity, so the technical approach must provide sufficient resolution and field of view to capture both levels. 3. Since movement preparation is a sub-second process, the neurophysiological recordings must also offer adequate time resolution to detect these neurophysiological variations. In this study, as illustrated in Figure 1, we sought to re-investigate these questions using functional ultrasound (fUSi), a recent imaging technique that provides both high temporal and spatial resolution images of Cerebral Blood Volume (CBV) changes^18^. In addition to the analysis of target Regions of Interest (ROIs), we used a simple Convolutional Neural Network (CNN) architecture to identify the brain state of a given fUSi frame, gaining insight into the global CBV pattern dynamics without spatial *a priori*. Our approach enabled us to extract unsupervised brain activity during single self-initiated action without the need for multilevel statistics, averaging, or filtering methods.

**Fig 1.**
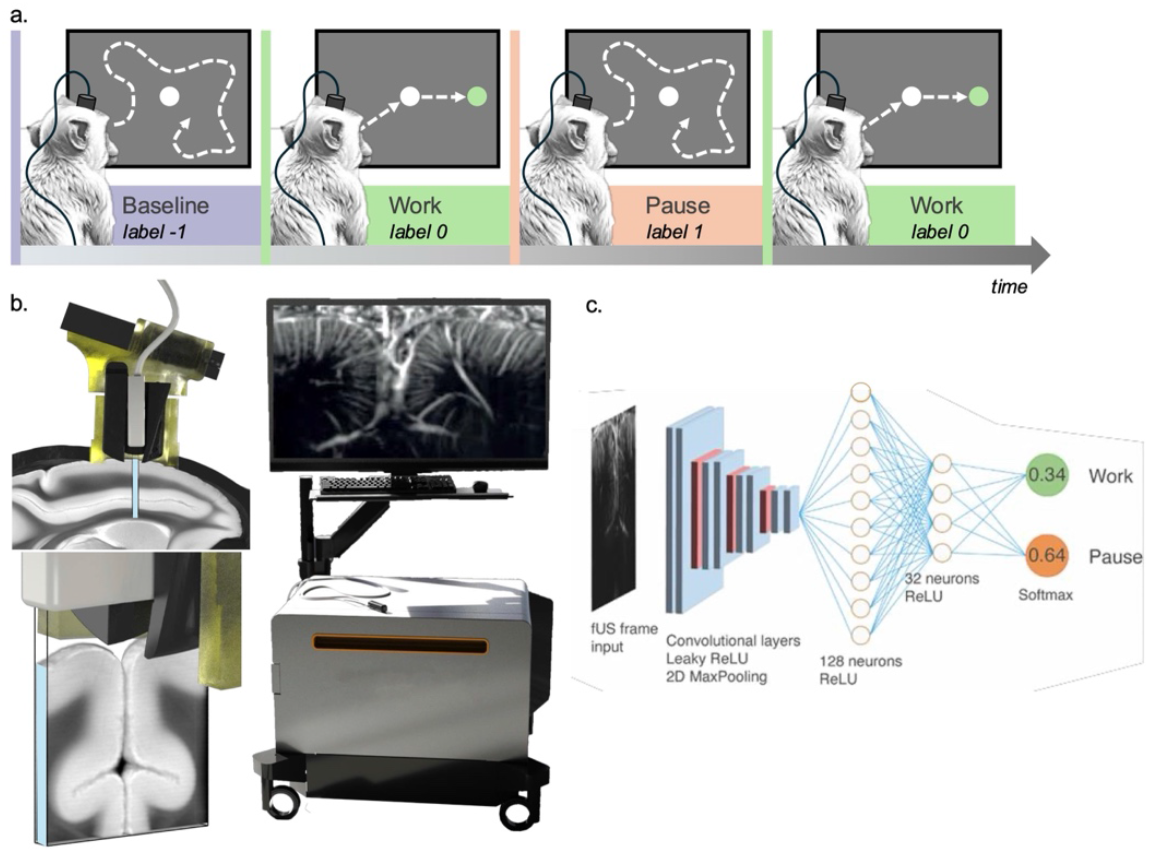
Experimental protocol, data labeling, and neural network architecture. (a) Oculomotor saccade task: Protocol consists of alternating work and pause periods. Animals are not engaged in the task during the resting state (purple, 200-220 seconds, random duration). Following this, a fixation point is presented (green timeline), followed by a cue indicating a saccade or antisaccade movement. The sequence of fixation and saccade is designated as “work” (green timeline). After completing a series of trials, animals can self-initiate pauses (orange timeline). Upon resuming activity, a new work period begins (green timeline). (b) Brain activity is monitored through functional ultrasound (fUSi) imaging during the whole task by measuring the cerebral blood volume (CBV). A 15 MHz ultrasonic probe is inserted in the recording chamber (AP + 23 mm, ML + 2 mm) and connected to an ICONEUS ONE prototype to record brain activity. (c) Using behaviour, data are labeled into three categories: baseline (−1), work (0), and pause (1) for further analysis through convolutional neural network (CNN) (d) Architecture of the CNN: the CNN utilizes four successive convolutional layers with LeakyReLU activation (blue) followed by 2D MaxPooling layers (red). This is followed by two fully connected layers of 128 and 32 neurons with ReLU activation. The final layer has two output neurons, enabling the CNN to classify an input fUSi frame as either pause or activity. The sum of the output probabilities always equals 1.

## RESULTS

Two captive-born rhesus macaques were trained to perform a saccade/antisaccade oculomotor task while their brain activity was recorded using fUSi. Each session lasted approximately one hour, during which the monkeys were undisturbed and not subjected to any interventions by the experimental staff to stimulate their participation.

We take the advantage of high signal to noise ratio of fUSi to examine brain activity during some sessions while the monkeys stopped and resumed tasks after some delays. The moment and duration of these “pause and resume events” were unpredictable. A posteriori, the resume event after a pause can be classified as a “self-generated” (SG) resume of tasks, in contrast to the “induced” start of the task, which occurs at the first trial during a session (Fig 1.b). Throughout the imaging session, we used fUSi to monitor the MFC neuro-vascular activity and assess the variation of CBV in our imaging plan during these different induced, pause and resume tasks (Fig 1.a)

First, we conducted a seed-based analysis on the Supplementary Eye Fields (SEF), which is strongly implicated in oculomotor tasks and whose activity can be measured using fUSi, as indicated by Claron and colleagues (Dizeux et al. 2019; Claron et al. 2023). Additionally, we examined the medial cingulate cortex (MCC), known for its role in action supervision^19^. Our results showed an increase in CBV before the effective end of the pause (EEoP). All results are given as mean±s.e.m.

As illustrated in Figures 2b and 2e, in the SEF, an increase in the CBV is observed at +0.4s (±1.3s) and +0.8s (±1.0s) relative to task onset, respectively, for monkey S and G (p > 0.05, 1 sample t-test). Relative to the Effective End of Pause (EEoP), an increase of the CBV within SEF is observed at −6.5s(±1.0s) and −7.2s(±1.0s), respectively, for monkey S and G. (p < 0.001 compared to the “onset” condition for both monkeys).

**Fig 2.**
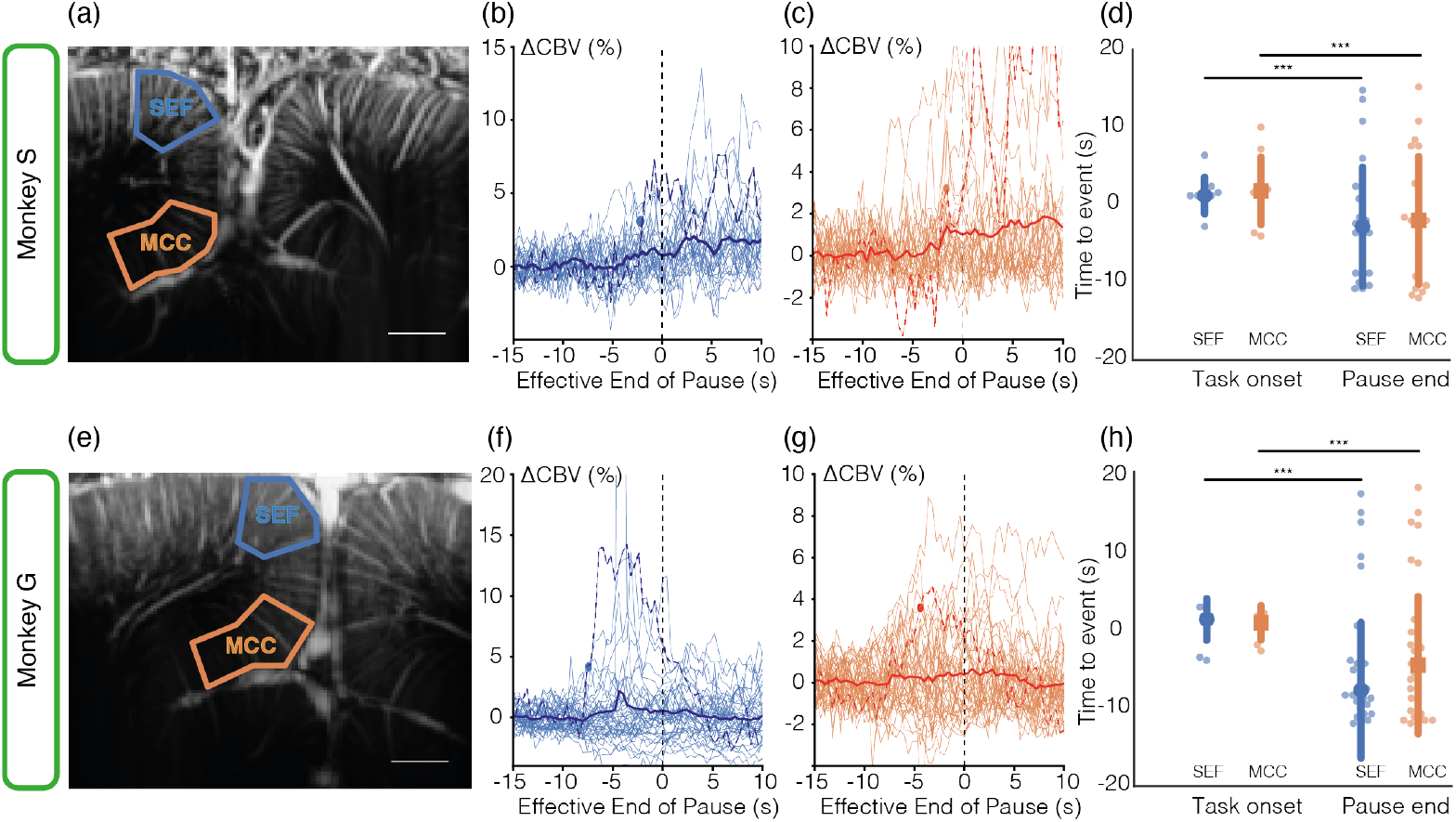
Region of Interest based onset analysis. (a) Anatomical map for Monkey S with 2 ROI: blue – SEF, orange – MCC, (b) Threshold (mean+1std over 2 timepoint) swarm plots for SEF for the Effective End of Pause (EEoP) for monkey S. For the EEoP, the threshold indicating ΔCBV increase is crossed −6.5±1.0s for the SEF and (c) −5.6±1.1s for the MCC. (d) Scatterplot of onset distribution for both ROI and both condition (*** p < 0.001). (e)-(h) same as (a)-(d) for monkey G. Thresholds: EEoP SEF: −7.2±1.0s, EEoP MCC: −5-5±1.2s.

As illustrated in Figures 2c and 2f, in the MCC, CBV increase occurred at 1.9s(±1.5s) and −0.6s(±1.0s) relative to task onset, respectively, for monkey S and G (p > 0.05, 1 sample t-test). Relative to the Effective End of Pause (EEoP) condition, an increase of the CBV within MCC is observed −5.6s(±1.1s) and −5.5s(±1.2s), respectively, for monkey S and G (p < 0.001 while compared to the “onset” condition for both monkeys)

As it was not depicted at the bold or vascular levels, these results provided a first description of the animals’ vascular and systemic action preparation within the fronto-medial cortex, which is essential for them to resume their tasks. However, the main limitation of this methodology was the strong assumptions made about the selected pixels we used and signed as SEF and the MCC ROIs. In a second step, we analyze the fUSi individual frames in their entirety without spatial *a priori*, we chose to train a convolutional neural network (CNN) for the classification of frames into pause or work states. Only the scar tissue on the top of the brain was excluded from the analysis.

The network architecture, as illustrated on Fig. 1.c, consisted of 3 convolutional layers and 2 fully connected layers before the softmax (sum of output neurons equal to 1) for classification into one of 2 labels: Pause / Work.

The network was trained independently for each imaging session using an equal number of pause and work frames, employing a stratified 5-folds cross-validation approach per session with randomly selected frames. Frames around state transitions, including those at the beginning and end of pauses, were excluded from the training set. The classification accuracy was 83.3% +/−9.77 and 90.3% +/−4.98 on average across acquisitions for subjects S and G respectively, and was statistically different from chance (i.e., a 50% accuracy on average), as confirmed by permutation tests with 1000 permutations. These results show that a single fUSi frame can contain enough information for the network to identify each individual’s behavioral state. This accuracy decreased by 29% and 35%, respectively, when evaluated on transition frames, this difference being statistically significant (p<0.001, t-test) in both cases. More specifically, the drop in accuracy was the smallest for transition frames located within a pause (start or end). It was significantly more prominent at the end of an activity period than at the start or end of pause periods. These drops suggest that modifying the spatial activity patterns in the brain is not immediate but somewhat transient or gradual. In particular, the fact that the spatial patterns learned by the network do not apply to the end of work- or pause-labelled periods indicates that these changes occur before the behavioral changes.

Subsequently, the trained network was applied to all the end-of-pause transition frames, providing each frame a probability of being labeled as either pause or work, which was normalized using the softmax operation. We observed that the network’s prediction accuracy dropped from 0.76±7.8.10^−3^ to 0.32±3.9.10^−3^, approximately 10 seconds before the actual end of the pause, transitioning from a probability of 1 (pause label) to 0 (work label) (cf. Fig 3.a). The prediction stabilized around 10 seconds after the effective end of the pause. At the effective end of pause (t=0), the prediction was 0.6±2.2.10^−3^, indicating nearly random guessing by the network. However, using the network output as our metric does not provide any insight into the critical areas of the imaging frame for classifying it as either pause or work.

**Figure 3:**
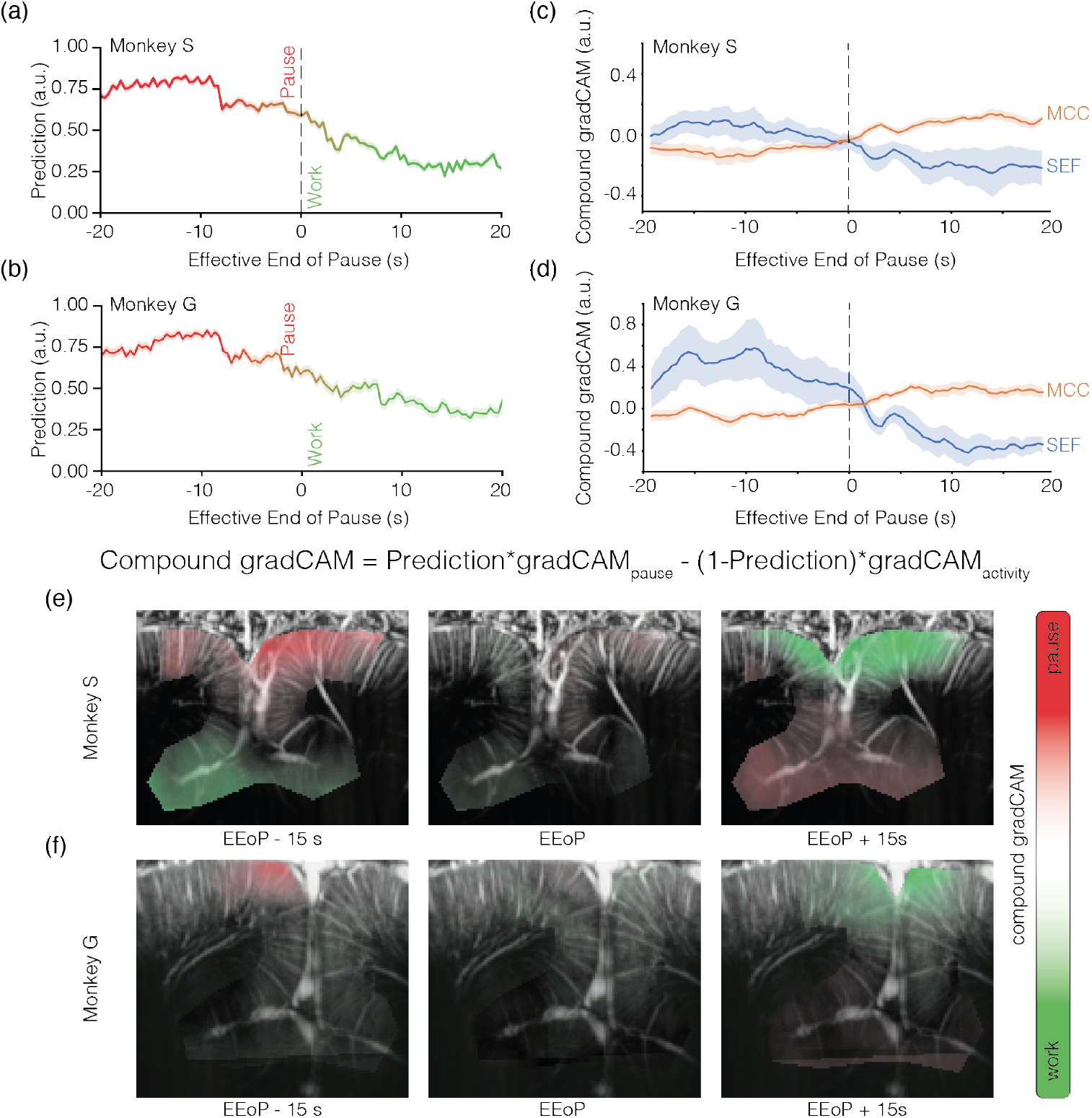
CNN prediction over transition frame, gradCAM++ output and compound gradCAM. (a) Prediction output over the transition frame for monkey S (b) same as (a) for monkey G (bottom) from −20s to 20s around the effective end of pause (EEoP). A value near 1 indicate a classification as pause, near 0, as work. mean+/−s.e.m (c) ROI average of the compound gradCAM for monkey S (top) and (d) same as (c) for monkey G (bottom). SEF in blue, MCC in orange, mean+/−std. Values are average z-score and show the importance of the area for the classification as either pause (positive values) or work (negative value). MCC seems to increase network instability whereas SEF seems to increase classification stability (e) Timepoint of the average compound gradCAM for monkey S and (f) monkey G over 3 timepoint (−15s, 0s, 15s). Red indicates an importance for classification as pause, green as work.

Using the gradCAM++ algorithm (Sup. fig. 1.), we were able to generate spatial importance maps (SIMs) highlighting for each frame the area’s most important for the classification, around the end of the pause (EEoP) and for each output label (pause or work). To visualize the brain regions most used by the network for each label, we normalized the SIMs into a z-score and created a linear combination of the pause and work SIMs based on the predicted pause output probability **α** using Eq.1.

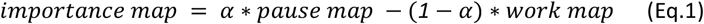

In the importance map of the monkey S, we observed a contrast between the importance of the SEF and MCC for the classification, as illustrated in Fig 3.b. Additionally, there was no significant important area around the EEoP. For monkey G., we observed the same behavior in the SIM. However, signals seemed to be weaker, especially in the MCC. This could be explained by the weaker signal-to-noise ratio in monkey G., possibly due to extra scar tissue and bone regrowth in the recording chamber. Altogether, the CNN and gradCAM++ results are consistent with the ROI based results, indicating our choice of analysis through compound gradCAM++ to be adapted to our analysis.

## DISCUSSION

We conducted a study to monitor neuronal activity in the MFC of two macaque monkeys during an oculomotor task. This region is known for producing and controlling purposeful eye movements in primates, particularly during visually guided saccades.

To tackle the question of the neurophysiological origin of voluntary preparation of movements, we benefited from a novel technology (fUSi) that permits a high signal-to-noise ratio from spatial changes in cerebrovascular blood volume (CBV) at the mesoscale level but at sufficiently fast time resolution to capture the concomitant variations of activity within MPFC.

We capture MPFC activity during breaks or the resumption of an ongoing voluntary single action. We compared this “time to resume act” to the initial trial induced into the session. Our findings indicated that activity in the supplementary eye field (SEF) and midcingulate cortex (MCC), both associated with eye movement, produced signals that effectively predicted the resumption of the task. This brain activity related to eye movement increased as the action was prepared and decreased during the time leading up to the pause period.

At first glance, these results align with clinical investigations that have demonstrated that damage to the MFC area of the brain in primates can lead to specific impairments in autonomic arousal when anticipating rewarded actions^5,20^. Among the MFC, observations in patients and quantifications in non-human primates with induced lesions indicate that the subgenual MCC is crucial for maintaining heightened autonomic arousal during reward anticipation and plays a vital role in regulating positive emotions^2,21,22^. However, the outcomes derived from uncontrolled extensive brain lesions in humans, or the methodologies used to induce lesions in animals are rather complex. As noted by the authors of these studies, it is possible that accidental damage to the white matter and beneath/above the subgenual MCC could also contribute to the observed effects. For instance, lesioning the subgenual MCC may involve transecting the corpus callosum and possibly damaging part of the dorsal part of MFC, including the SMA and PreSMA. This complexity makes it difficult to disentangle the specific contributions of the various cortical areas within this large region of the primates’ brain^3,23^. Keller and Heckhausen^24^ found an RP before involuntary movement and Haggard and Eimer^25^ and Hermann (2008) both found identical early RP before movements of either hand, suggesting the early RP is related to a more general process of anticipation and no to the preparation of a specific movement consistent with results from Trevena and Miller^26^. Pockett and Purdy^27^ reported voluntary movements without RPs. Schurger^28^ and Schmidt^29^ and Maoz^30^ have all demonstrated that the RP plausibly arises from averaging the random background noise or slow cortical potentials. Alexander^31^ found that RPs are also seen before decisions that do not involve movement. Verleger^32^ found that the onset of RPs depended strikingly on the interval between movements, and Maoz^30^ found no RPs before deliberate choice movements. These results make it unlikely that RP is a valid measure of genuine brain activity preparing for a voluntary movement. By using tone probes Hallet and Matsuhashi and Verbaarschot^33^ have shown that the onset of the intention to move occurs long before Libet’s W times and often before the onset of the RPs At a closer look, our results permit significant strides in understanding the neural mechanisms that facilitate movement; accurately identifying the moment conscious intention occurs remains a complex and elusive challenge. On a methodological level, our findings underscore the efficacy of employing CBV change measurements through fUSi. This innovative technique can effectively identify the neural signatures associated with motor control^18^. It offers new insights into the delicate interplay between gaze shifting—an essential aspect of visual attention—and the cognitive mechanisms responsible for maintaining repetitive efforts in tasks that require sustained attention and physical coordination.

Our findings further demonstrate that the intention to act can begin several seconds before an animal executes an action. This concept is not novel. In 1983, Libet’s experiments revealed a complex relationship between intention and action, igniting extensive debates across various fields, including neuroscience, philosophy, and law, particularly concerning free will and moral responsibility. Our data reveals that these neural processes are influenced by extensive signaling within the MFC, a key area involved in planning and executing movements.

Our data analysis indicated a switching of activity patterns between two critical brain regions: the SEF and the MCC. This dynamic of interactions between these regions suggests a possible concurrent physiological mechanism for generating voluntary actions. This early activity could be viewed not in terms of unconscious decision processes but rather by a process in which a decision is informed. Intention consciousness does not appear instantaneously but builds up progressively, and early neural markers of decision outcomes are not unconscious but reflect conscious goal evaluation stages, which are not final yet and, therefore, not reported with the Libet method.

## METHODS

### Animal model and behavioral data

All experiments were ethically approved by the French “Ministère de l’Education, de l’Enseignement Supérieur et de la Recherche” under the project reference APAFIS #6355-2016080911065046. Functional data were acquired from two captive-born macaques (Macaca mulatta), S and G, trained to perform various kinds of visual tasks. In the saccade task, the animal has to fix its gaze on the cue object presented on the right or left side of the screen; in the antisaccade task, it has to fix its gaze on the opposite side from where the cue appeared. Each animal performed a baseline (random duration from 200s to 220s) followed by saccades and antisaccades (randomized) over 1 hour. During data acquisition, the eye position of the primate was monitored at 1 kHz with an infrared video eye tracker (Eyelink 1k, SR-Research), which enabled live control of the behavioral paradigm and the delivery of a reward based on the success or failure of a visual task^34^.

#### Implant and probe for functional ultrasound imaging in awake behaving monkeys

The monkey head was fixed using a surgically implanted titanium headpost (Crist Instrument, MD, USA). After behavioral training of the animals, a recording chamber (CILUX chamber, Crist Instrument, MD, USA) was implanted and a craniotomy (diameter 19 mm) was performed (mediolateral +0mm, anteroposterior : +26mm). A custom ultrasonic probe (128 elements, 15 MHz, 100 × 100 µm^2^ of spatial resolution) with ultrasonic gel was used in the chamber. The acquired images had a pixel size of 100 × 100 µm and a slice thickness of 400 µm.

### Functional ultrasound (fUSi) recording

Cerebral blood volume (CBV) variations were measured with an ultrasonic imager (Iconeus, Paris, France). Data were acquired by emitting continuous groups of 11 planar ultrasonic waves tilted at angles varying from −10° to 10°. Ultrasonics echoes were summed to create a single compound image acquired every 2ms. After a spatiotemporal filtering based on the singular value decomposition of these ultrasonic images, final Doppler images were created by averaging 200 compound ultrasonic images. N = 19 sessions and N = 10 sessions were used for monkey S and G respectively.

### Eye movements Pupil recordings

During tasks, eye movements and pupil diameter were recorded using a video eye tracker (Eyelink 1k, SR-Research) connected to an analog-to-digital converter (Plexon Inc, TX, USA). All datas were collected on Plexon software and analyzed on MATLAB (The MathWorks Inc., Massachusetts, USA).

*fUSi data labeling*

### Behavioral labeling

During a session, the fUSi was recorded for around 75 min. The start of the recording consist on a grey screen (#3F3F3F) on which nothing is displayed for 200 to 220 seconds (random). Trials start after this baseline, consisting of a fixation followed by a saccade or an anti-saccade. Trials are separated by a intertrial (grey screen, #3F3F3F) of 4 to 5 seconds (random). If the animal does not perform the fixation in the following 1s after the proposition, intertrial (4 to 5 seconds) starts.

### Pauses

A pause is defined by:

- Start: five fixations proposed but not realized. The 1st fixation of the consecutive five is the start of the pause.
- End: At least one fixation followed by a saccade or an anti-saccade is enough to end the pause

This allowed the labeling of data for CNN training. The labels were chosen: −1 correspond to baseline, 0 to work, 1 to pause.

However, during further analysis on ΔCBV and predictions, only the pause endings where the animal was in pause for at least 20 seconds before and worked for at least 20 seconds after the end were kept for analysis.

### Region of Interest-based Analysis

#### fUSi signal extraction

fUSi signals are extracted by manually drawing a region of interest over the chosen area (SEF and MCC) before a spatial non-weighted average over the ROI. After spatial extraction, the signal was temporally cut using the above-mentioned criteria for pause endings, then normalized into local ΔCBV using the following expression:

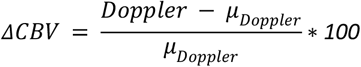

where μ_Doppler_ is the average of the local signal over the 20 first points (corresponding to −20s to −12s)

#### Threshold analysis

To analyze the increase of ΔCBV about t at the end of the pauses, we are looking for at least 2 consecutive time points where ΔCBV(t) exceeds the mean (ΔCBV_[-20:-12]_)+std(ΔCBV_[-20:-12]_) from −20 to −12 seconds, plus one standard deviation of ΔCBV over the same interval. The first of these time points will be designated as the threshold crossing time. If no such point is identified within the interval of −20 to 20 seconds, that point will be excluded from further analysis.

Additionally, we will assess the normality of the data distribution using the Shapiro-Wilk test before applying a Welch’s test to account for heteroscedasticity between the “task onset” and “pause” groups.

### Convolutional Neural Network analysis

#### Data preparation

fUSi frames were prepared by manually masking out the outside of the brain (corresponding pixels were set to 0) for all frames of each acquisition. Each frame was then divided pixel-by-pixel by the corresponding average value in the baseline. A correction to the global intensity drift observable in each acquisition was then applied pixel-wise by removing from the temporal data a numeric component, corresponding to a second-degree polynomial fit of the image intensity variation across the acquisition. The dataset for each acquisition was built using all pause frames and the same number of work frames selected randomly across the acquisition (except in the baseline). Transitions corresponding to 50 frames before and after each behavioral state change were excluded from the training at this stage. A total of 80% of frames were randomly chosen within this balanced dataset to build the training and validation sets, and the remaining 20% were set aside for accuracy testing.

#### Artificial neural network architecture

A convolutional neural network, built using the Keras library and TensorFlow environment, was trained to classify fUSi images into work and pause frames, labeled as explained above. The neural network took as input the full 128×112 pixel fUSi image and consisted of 4 consecutive convolution layers with 16, 32, 64, and 128 filters, respectively, a kernel size of 3×3 initialized with He initialization, a stride of 1 in each direction with « same » padding, and ReLU activation. Each convolutional layer underwent batch normalization and was followed by a 2D MaxPooling layer with a pool size of 2×2 and a stride of 2 in each direction. These were followed by two dense layers of 128 and 32 neurons with ReLU activation, HeUniform kernel initialization and a 15% dropout rate. The output was a two-neuron softmax layer with Glorot Uniform initialization, corresponding to the 2 classes of work and pause. Acquisitions, where the ratio of work to pause frames did not allow for a balanced training/validation dataset of more than 500 frames, were removed from the analysis.

#### Training

For each acquisition, training was done using Adam optimizer for a maximum of 1500 epochs. Early stopping was used when the validation accuracy varied by less than 0.001 for more than 50 epochs, to prevent potential overfitting. The best weights were restored after stopping. The learning rate was set at 10^−4^ and was reduced by a factor of 0.2 when reaching a plateau of the validation accuracy within a delta of 0.01 for at least 10 epochs, down to a minimum of 10^−6^. The training was done on the training/validation set by 5-fold cross-validation using the StratifiedKFold function of the sci-kit-learn library for balanced folds.

* To avoid potential information from one frame to the next, all test frames were separated by at least one sample frame from the closest training frame in the acquisition. *

#### Model evaluation

Each model (i.e. each fold for each acquisition) was evaluated by calculating the percentage of test cases for which the model classification was correct. The network classification was obtained by selecting the neuron with the highest probability output, i.e. work or pause. Accuracy evaluation was done on the original test dataset, including only frames not located within 50 frames of a transition, as well as on the « transition frames », i.e. all frames within 50 frames of a transition event. The comparison between accuracy values obtained for different groups of frames was done with statistical testing using the T-test when the distributions of values were normal (according to the Shapiro-Wilk test with a p-value cutoff of 0.05) and the Mann Whitney-U test otherwise.

#### Model prediction at transitions

Since the accuracy at transitions dramatically dropped compared to the accuracy calculated on non-transition frames, we aimed to display the typical accuracy around a transition event from pause to work. This was done by accumulating the predicted network output, i.e. the probability that a given frame corresponds to the pause state, for all models and all corresponding transitions. Transitions between pause and work periods shorter than 100 frames were excluded from the analysis. For each time point before the transition, we calculated the statistical difference between the output values of the model before this time point and the output of the model 15 frames after the transition using the Wilcoxon signed-rank test. This was compared to the output of a simple threshold model predicting the behavioral state based on the average value of the MCC and SEF regions. This model was designed by selecting the best among three threshold values corresponding to M+n*sigma where M is the average pixel value, sigma is the standard deviation of values and n a parameter varying from 1 to 3.

#### Spatial and temporal importance maps

Importance maps were generated for each fUSi frame within the preparation period (50 pause frames before work restart) using the gradCAM++^35^ technique for the target class of work or pause corresponding to the frame label. These maps were averaged for each transition and each acquisition. In addition, for each transition, we identified the time point before the transition at which each pixel reached its maximal importance gradCAM++ for the classification. Only regions reaching gradCAM++ scores above 0.2 were considered. This result was then reported, averaged on all transitions for each acquisition, to provide an insight into the spatio-temporal activation patterns at transitions.

#### gradCAM++ map compound

All Doppler maps are recalibrated in the same space using a combination of AFNI and FSL though Nipype warper Python library. Doppler data are transformed into NIFTI files using the Nibabel python package and transformed back into matrices using the same package. The NIFTI header is the base one from Nibabel. N = 12 and N = 4 sessions were used for monkey S and G. Lost sessions were lost because of the recalibration in the common space that was not possible for all of them.

The gradCAM++ maps were generated for each timepoint through acquisition for each output, cut using the same condition as defined before, transformed into z-score and combined as :

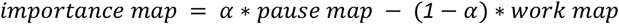

with *α* being the pause probability prediction given as output by the full network (1 : pause, 0 : work)

## Acknowledgments

JC was funded by FRM postdoctoral fellowship (SPF202110014064). ANR #. PHYSMED

## FIGURES

**Supplementary Figure 1:**
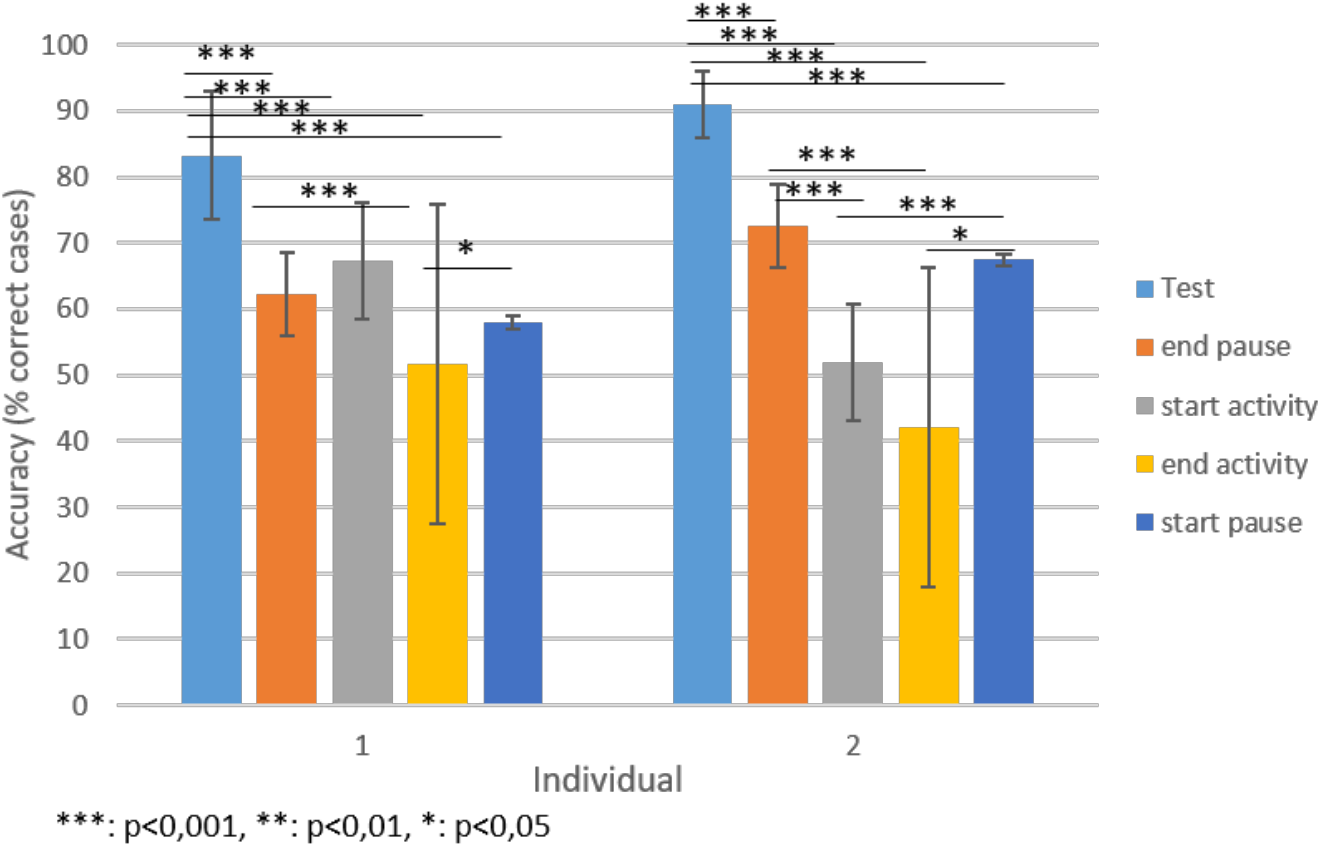
Classification accuracy of the neural networks for each individual. **Accuracy is given as the percentage of correctly classified test cases, averaged across network realizations and acquisitions for each individual, with error bars corresponding to the standard deviation. Also shown are the accuracy at transitions at the end of a pause, at the start of an activity period, at the end of an activity period, and at the start of a pause.**

**Supplementary Figure 2:**
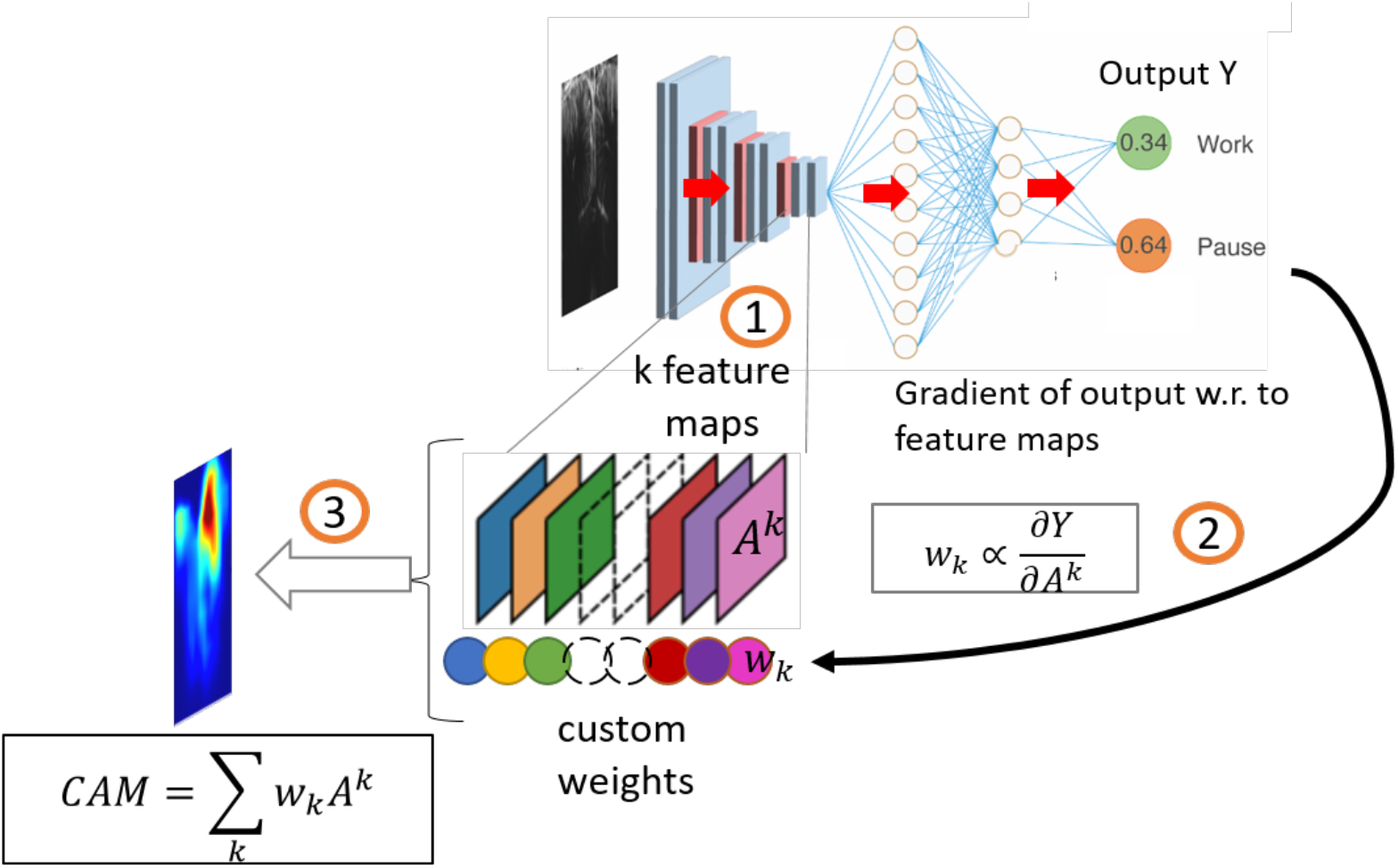
Method / Network classification & map of importance. Illustration of the gradCAM++ method to build class activation maps (CAM): 1. The image goes through the classification network, and the feature maps achieved in the last convolution layer are extracted. 2. The output of the network for the considered class is calculated, and the gradient of that output with respect to the last layer feature maps is used to define a custom weight. 3. The CAM is built as a linear combination of the last layer feature maps with the weights calculated in 2.

## REFERENCES

1. Libet, B., Gleason, C. A., Wright, E. W. & Pearl, D. K. Time of conscious intention to act in relation to onset of cerebral activity (readiness-potential). The unconscious initiation of a freely voluntary act. Brain J. Neurol. 106 (Pt 3), 623–642 (1983).

2. Monosov, I. E., Haber, S. N., Leuthardt, E. C. & Jezzini, A. Anterior Cingulate Cortex and the Control of Dynamic Behavior in Primates. Curr. Biol. CB 30, R1442–R1454 (2020).

3. Oerlemans, J. et al. Unravelling the origin of reward positivity: a human intracranial event-related brain potential study. Brain 148, 199–211 (2025).

4. Lau, H. C., Rogers, R. D., Haggard, P. & Passingham, R. E. Attention to intention. Science 303, 1208–1210 (2004).

5. Haggard, P. Human volition: towards a neuroscience of will. Nat. Rev. Neurosci. 9, 934–946 (2008).

6. Desmurget, M. et al. Movement intention after parietal cortex stimulation in humans. Science 324, 811–813 (2009).

7. Fried, I., Mukamel, R. & Kreiman, G. Internally generated preactivation of single neurons in human medial frontal cortex predicts volition. Neuron 69, 548–562 (2011).

8. Hallett, M. Physiology of free will. Ann. Neurol. 80, 5–12 (2016).

9. Schultze-Kraft, M. et al. Suppress Me if You Can: Neurofeedback of the Readiness Potential. eNeuro 8, ENEURO.0425-20.2020 (2021).

10. Seghezzi, S., Zirone, E., Paulesu, E. & Zapparoli, L. The Brain in (Willed) Action: A Meta-Analytical Comparison of Imaging Studies on Motor Intentionality and Sense of Agency. Front. Psychol. 10, 804 (2019).

11. Cui, H. & Andersen, R. A. Posterior parietal cortex encodes autonomously selected motor plans. Neuron 56, 552–559 (2007).

12. Haggard, P. The Neurocognitive Bases of Human Volition. Annu. Rev. Psychol. 70, 9–28 (2019).

13. Pesaran, B., Nelson, M. J. & Andersen, R. A. Free choice activates a decision circuit between frontal and parietal cortex. Nature 453, 406–409 (2008).

14. Zapparoli, L., Seghezzi, S. & Paulesu, E. The What, the When, and the Whether of Intentional Action in the Brain: A Meta-Analytical Review. Front. Hum. Neurosci. 11, 238 (2017).

15. Zapparoli, L. et al. Dissecting the neurofunctional bases of intentional action. Proc. Natl. Acad. Sci. 115, 7440–7445 (2018).

16. Ariani, G., Wurm, M. F. & Lingnau, A. Decoding Internally and Externally Driven Movement Plans. J. Neurosci. Off. J. Soc. Neurosci. 35, 14160–14171 (2015).

17. Paus, T. Primate anterior cingulate cortex: where motor control, drive and cognition interface. Nat. Rev. Neurosci. 2, 417–424 (2001).

18. Macé, E. et al. Functional ultrasound imaging of the brain. Nat. Methods 8, 662–664 (2011).

19. Stoll, F. M. et al. The Effects of Cognitive Control and Time on Frontal Beta Oscillations. Cereb. Cortex N. Y. N 1991 26, 1715–1732 (2016).

20. Darby, R. R., Joutsa, J., Burke, M. J. & Fox, M. D. Lesion network localization of free will. Proc. Natl. Acad. Sci. 115, 10792–10797 (2018).

21. Vázquez, D. et al. Anterior cingulate cortex lesions impair multiple facets of task engagement not mediated by dorsomedial striatum neuron firing. Cereb. Cortex N. Y. N 1991 34, bhae332 (2024).

22. Li, Y. S., Nassar, M. R., Kable, J. W. & Gold, J. I. Individual Neurons in the Cingulate Cortex Encode Action Monitoring, Not Selection, during Adaptive Decision-Making. J. Neurosci. 39, 6668–6683 (2019).

23. Holroyd, C. B. & Umemoto, A. The research domain criteria framework: The case for anterior cingulate cortex. Neurosci. Biobehav. Rev. 71, 418–443 (2016).

24. Keller, I. & Heckhausen, H. Readiness potentials preceding spontaneous motor acts: Voluntary vs. involuntary control. Electroencephalogr. Clin. Neurophysiol. 76, 351–361 (1990).

25. Haggard, P. & Eimer, M. On the relation between brain potentials and the awareness of voluntary movements. Exp. Brain Res. 126, 128–133 (1999).

26. Trevena, J. A. & Miller, J. Cortical movement preparation before and after a conscious decision to move. Conscious. Cogn. Int. J. 11, 162–190 (2002).

27. Pockett, S. & Purdy, S. C. Are Voluntary Movements Initiated Preconsciously? The Relationships between Readiness Potentials, Urges, and Decisions. in Conscious Will and Responsibility: A Tribute to Benjamin Libet (eds. Sinnott-Armstrong, W. & Nadel, L.) 0 (Oxford University Press, 2010). doi:10.1093/acprof:oso/9780195381641.003.0005.

28. Schurger, A., Sitt, J. D. & Dehaene, S. An accumulator model for spontaneous neural activity prior to self-initiated movement. Proc. Natl. Acad. Sci. U. S. A. 109, E2904–2913 (2012).

29. Schmidt, S., Jo, H.-G., Wittmann, M. & Hinterberger, T. ‘Catching the waves’ - slow cortical potentials as moderator of voluntary action. Neurosci. Biobehav. Rev. 68, 639–650 (2016).

30. Maoz, U., Yaffe, G., Koch, C. & Mudrik, L. Neural precursors of decisions that matter—an ERP study of deliberate and arbitrary choice. eLife 8, e39787 (2019).

31. Alexander, P. et al. Readiness potentials driven by non-motoric processes. Conscious. Cogn. 39, 38–47 (2016).

32. Verleger, R., Grauhan, N. & Śmigasiewicz, K. Go and no-go P3 with rare and frequent stimuli in oddball tasks: A study comparing key-pressing with counting. Int. J. Psychophysiol. 110, 128–136 (2016).

33. Verbaarschot, C., Haselager, P. & Farquhar, J. Detecting traces of consciousness in the process of intending to act. Exp. Brain Res. 234, 1945–1956 (2016).

34. Valero-Cabre, A. et al. Frontal Non-Invasive Neurostimulation Modulates Antisaccade Preparation in Non-Human Primates. PLOS ONE 7, e38674 (2012).

35. Chattopadhay, A., Sarkar, A., Howlader, P. & Balasubramanian, V. N. Grad-CAM++: Generalized Gradient-Based Visual Explanations for Deep Convolutional Networks. in 2018 IEEE Winter Conference on Applications of Computer Vision (WACV) 839–847 (2018). doi:10.1109/WACV.2018.00097.

